# A quantitative description of the mouse piriform cortex

**DOI:** 10.1101/099002

**Authors:** Shyam Srinivasan, Charles F Stevens

## Abstract

In this report, we provide a quantitative description for 2 aspects of the mouse piriform cortex. First, we give volumetric estimates of regions within the anterior and posterior piriform cortex. Second, we estimate the neuronal densities in each of these regions. From these two estimates, we calculate that the mouse piriform contains around half a million neurons equally distributed over the anterior and posterior piriform cortex. Quantitative descriptions such as these are important because they make it possible to construct realistic models and provide a constraint that theories of the olfactory circuit must fulfil. We show how quantitative descriptions can be useful for modelling by using our data to refine and improve earlier models of piriform cortex activity.

## Background

In this report, we provide a quantitative description of the mouse piriform cortex (Neville & Haberly 2004; Bekkers & Suzuki 2013). The piriform cortex, also called the pyriform or pre-piriform cortex, is the largest olfactory region in the cortex, and is responsible for encoding experience and learning dependent odors. It forms the third stage of the olfactory circuit that begins with the olfactory epithelium in the nose followed by the olfactory bulb. A quantitative description of the piriform cortex is needed for 2 reasons.

First, the organization of the piriform cortex differs from other sensory cortices like the auditory or visual cortex which are stereotypically organized (Hubel & Wiesel 1962). In the piriform cortex, odors seem to be coded by a random and sparse ensemble that does not seem to follow an obvious stereotypic organization (Stettler & Axel 2009; Illig & Haberly 2003; Rennaker et al. 2007; Poo & Isaacson 2009). For that reason, theories from sensory cortices that are stereotypically organized cannot be applied to the piriform. Studies have quantitatively described the first two stages in the olfactory epithelium and the bulb (Murthy 2011; Soucy et al. 2009; Kawagishi et al. 2014; Royet et al. 1988; Richard et al. 2010). Around 5 million OSNs communicate odor information to around 1800 or 3700 glomeruli in the olfactory bulb. In order to develop a theory, we need quantitative information of the third stage.

Second, theoretical studies have linked visual cortex function to its anatomy by using the theory of scaling (Srinivasan et al. 2015; Stevens 2001). These relate how visual cortex function will improve with an increase in cortex size. Such predications cannot be made for the olfactory circuit, because numbers for the piriform cortex are lacking. This report takes a step in addressing the gap in such quantitative descriptions. We, next provide a summary of the piriform cortex components that were estimated.

## Summary

The piriform cortex is tri-laminar and a part of the paleocortex (Neville & Haberly 2004). It is located caudal to the bulb, and abuts the anterior olfactory nucleus as shown in Figure 1. Morphologically, the piriform is easily distinguishable in coronal sections because of the extreme density of cell bodies in its Layer II. We measured two aspects of piriform cortex morphology. First, we measured the volumetric aspect, which includes measurements of layer widths, the surface area and volume of the piriform cortex. Second, we measured the number of neurons in each of layer under a square mm of cortical surface (both measures are described in the next section; see Figures 1, 2, and methods). From these two measures in 3 mice, we estimated that the piriform cortex, on average, contains 532,000 neurons. As all these measures were for one hemisphere, the total number of piriform neurons in the mouse is about a million.

**Figure 1:**
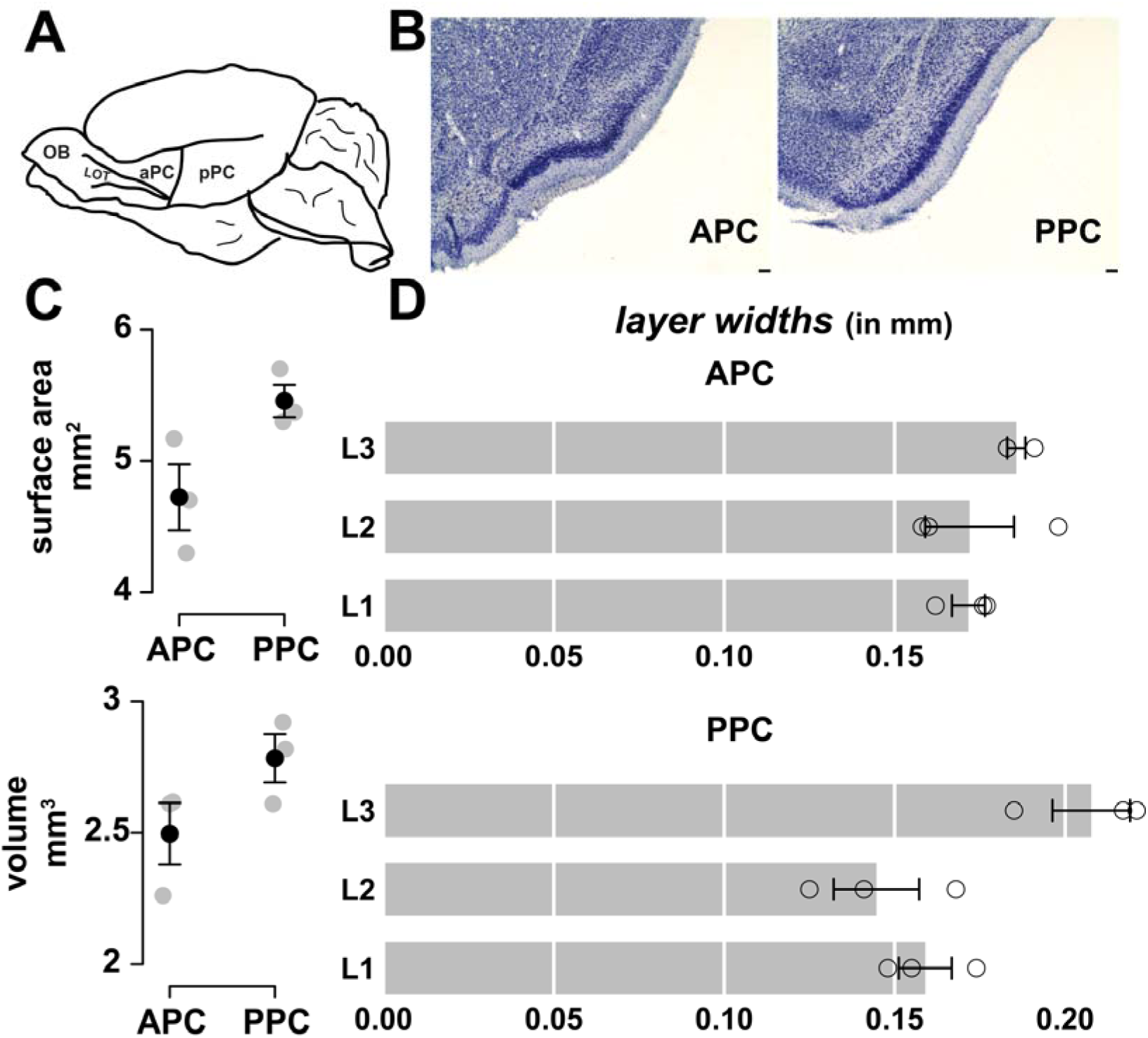
Volume and Surface area estimates of the mouse piriform cortex. (A) Schematic of the mouse brain adapted from (Bekkers & Suzuki 2013), by tracing the outline of the mouse brain. (B) Representative sections of the mouse piriform cortex in the anterior and posterior. The anterior piriform cortex is distinguished from the posterior by the presence of the lateral olfactory tract. Scale Bar: 50 μm. (C) top panel: Surface area of the piriform cortex in the anterior (4.72 +/− 0.25 mm^2^) and posterior (5.45 +/− 0.12 mm^2^), bottom panel: Volume of the anterior (2.49 +/− 0.11 mm^3^) and posterior piriform cortex (2.78 +/− 0.09 mm^3^). (D) Widths of Layers 1, 2, and 3 in the anterior and posterior piriform cortex. Layer 1:0.171 +/−.004 mm and 0.159 +/−.007 mm, Layer 2:0.172 +/− .013 mm and 0.144 +/−.012 mm, Layer 3:0.185 +/−.002 mm and 0.207 +/−.011 mm. Referto Tables 3-5 for measurements in individual mice. Error bars and measures are standard error.

**Figure 2:**
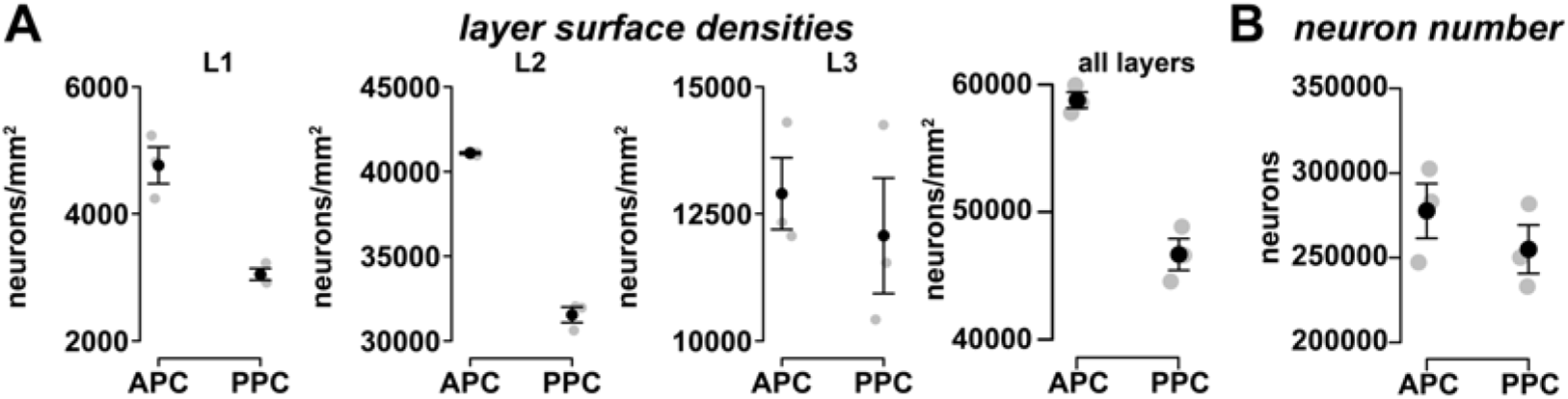
The mouse piriform cortex contains half a million neurons. (A) Surface area neuronal densities (number of neurons under 1 square mm of the cortex) of the piriform cortex for layers 1, 2, and 3, and all layers combined. Layer 1: 4,765 +/− 287 (APC) and 3,048 +/− 92 (PPC), Layer 2: 41,089 +/− 69 (APC) and 31,538 +/− 460 (PPC), Layer 3:12,899 +/− 706 (APC) and 12,070 +/− 1,137 (PPC), All layers: 58,753 +/− 801 (APC) and 46,655 +/− 1,583 (PPC) (B) Total number of neurons in the mouse anterior piriform cortex (APC) and PPC: 277,643 +/− 16,208 and 254,948 +/− 14,303. Refer to Tables 1, 2, and 6 for measurements in individual mice. Error bars and measures are with standard error.

## A quantitative description of the piriform cortex

In this section, we provide a quantitative description of the piriform cortex. The piriform cortex can be morphologically and functionally divided into the anterior piriform cortex (APC) and the posterior piriform cortex (PPC). Morphologically, the APC is distinguished by the presence of the lateral olfactory tract (Neville & Haberly 2004). Functionally, while activity in the APC has been shown to distinguish amongst odor objects (Stettler & Axel 2009; Rennaker et al. 2007), the posterior has been shown to distinguish odor quality (Gottfried 2010). For this reason, we present our measurements of the APC and PPC separately.

First, our volumetric measurements showed that, on average, the piriform cortex has a volume of 2.49 mm^3^ in the anterior and 2.78 mm^3^ in the posterior (this and subsequent measures are averages from 3 mice, Figure 1C). Correspondingly, its surface area in the anterior is 4.72 mm^2^ and 5.45 mm^2^ (Figure 1C, Table 5). Our measurements also showed that Layer 3 was consistently the thickest layer having a width of 0.185 mm in the anterior, and 0.207 mm in the posterior (Figure 1D, Tables 3, 4). Layers 1 and 2 were thinner with widths of 0.17 mm and 0.17 mm in the anterior, and 0.159 and 0.146 mm in the posterior.

**Table 1.** Neuron density (per sq. mm of surface area) in each layer of the APC (3 mice). For all tables, each box gives the mean for that mouse and the standard error of the measurements.

|  | Mouse 1 | Mouse 2 | Mouse 3 |
| --- | --- | --- | --- |
| Layer 1 | 4,819;414 | 5,235;629 | 4,242;617 |
| Layer 2 | 41,119; 2,581 | 40,957;3,106 | 41,191;2,104 |
| Layer 3 | 14,304;1,000 | 12,331;1,142 | 12,062;1,793 |

**Table 2.** Neuron density (per sq. mm of surface area) in each layer of the PPC

|  | Mouse 1 | Mouse 2 | Mouse 3 |
| --- | --- | --- | --- |
| Layer 1 | 3,226;1,004 | 3,003;419 | 2,916;560 |
| Layer 2 | 31,960;2,543 | 32,036;3,026 | 30,619;2,628 |
| Layer 3 | 14,252;1,218 | 11,538;1,304 | 10,420;1,017 |

**Table 3.** Widths of ayers in the APC (in mm).

|  | Mouse 1 | Mouse 2 | Mouse 3 |
| --- | --- | --- | --- |
| Layer 1 | .176;.006 | .162;.015 | .177;.005 |
| Layer 2 | .198;.01 | .160;.009 | .158;.002 |
| Layer 3 | .183;.002 | .183;.015 | .191;.009 |

**Table 4.**
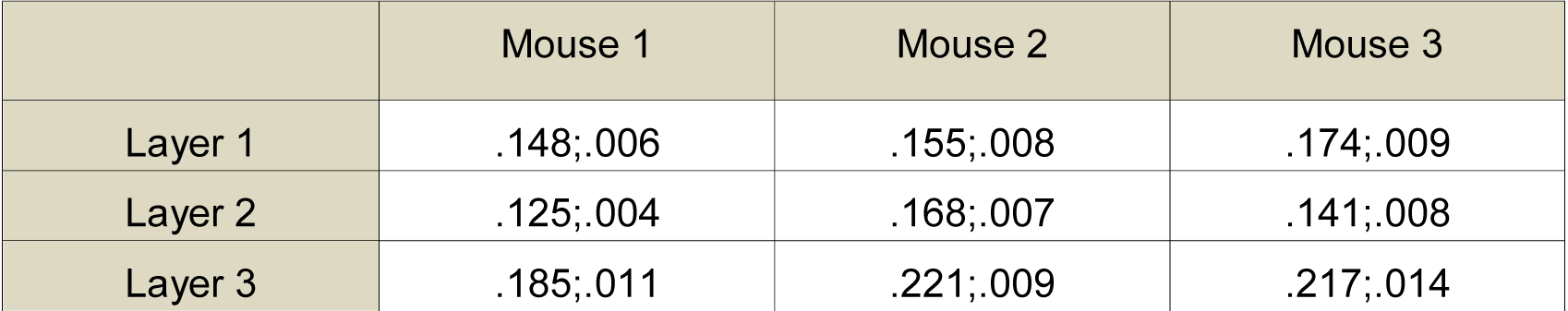
Widths of layers in the PPC (in mm)

|  | Mouse 1 | Mouse 2 | Mouse 3 |
| --- | --- | --- | --- |
| Layer 1 | .148;.006 | .155;.008 | .174;.009 |
| Layer 2 | .125;.004 | .168;.007 | .141;.008 |
| Layer 3 | .185;.011 | .221;.009 | .217;.014 |

**Table 5.** Surface area measurements (in mm^2^)

|  | APC | PPC | Entire PC |
| --- | --- | --- | --- |
| Mouse 1 | 4.7 | 5.7 | 10.4 |
| Mouse 2 | 5.17 | 5.37 | 10.5 |
| Mouse 3 | 4.3 | 5.3 | 9.6 |

Second, we found that the surface area density of neurons under a square mm was 58,753 neurons/mm^2^ in the anterior and 46,655 neurons/mm^2^ in the posterior (Figure 2A, Tables 1, 2). These neurons were distributed over Layers 1, 2, and 3, which had 4,765, 41,089, and 12,899 neurons/mm^2^ in the APC, and 3,048, 31,538, and 12,070 neurons/mm^2^ in the PPC (Figure 2A, Tables 1, 2). Combining these estimates with estimates of surface area, we calculated the total number of neurons to be 278,000 in the anterior, and 255,000 in the posterior (Figure 2B, Tables 6).

**Table 6.** All layers, number of neurons, *Total numbers of neurons*

|  | APC | PPC |
| --- | --- | --- |
| Mouse 1 | 283,187 | 281,796 |
| Mouse 2 | 302,563 | 250,118 |
| Mouse 3 | 247,228 | 232,961 |

## Refining previous models of the primary olfactory cortex with quantitative data

We now illustrate how these estimates can be used to improve existing quantitative models representing the 2^nd^ and 3^rd^ stages of the olfactory circuit. An elegant study which showed that odors are encoded by a sparse and seemingly random ensemble of piriform neurons also presented a model explaining how such activity could arise (Stettler & Axel 2009). Although the model captured the essential architectural features of the olfactory circuit, the absence of quantitative data for the piriform cortex hindered its efficacy. A subsequent study of the analogous circuit in flies (Stevens 2015) with the same architectural logic overcame this issue because of the availability of quantitative data describing the 3^rd^ stage of the circuit as well as the connection characteristics between the 2^nd^ and 3^rd^ stages (Caron et al. 2013; Gruntman & Turner 2013). Our report, by providing similar numbers for the mouse circuit, makes similar realistic models, possible.

The main component in both models (Stettler & Axel 2009; Stevens 2015) was the connectivity matrix between ‘g’ glomeruli and ‘n’ 3^rd^ stage neurons. If the activity pattern of glomeruli were represented by a glomerular vector G, the activity vector of the 3^rd^ stage neurons in a linear spike rate coding model would be, *N* = *Conn. G*, where Conn is a matrix describing the connections from glomeruli to 3^rd^ stage neurons.

This model requires 3 quantities before one can use it to develop theories of olfaction: the number of glomeruli, the number of neurons, and the strength of synaptic connection between any glomerulus *‘i’* and neuron *‘j’*, i.e. the entry (*i*,*j*) in the connection matrix. The number of glomeruli is already known, and the number of neurons is available from this report. To complete the model, therefore, we need an estimate of the synaptic connection strength.

In a seemingly random network such as the one between glomeruli and piriform neurons, the synaptic connection strength between any glomerulus-neuron pair (*i*,*j*) can be obtained by sampling from random distributions that can accurately describe two components of connectivity. The first component is the number of synapses between pair (*i*,*j*). As studies have shown that synapses made by a particular glomerulus have no spatial preference, the number of synapses between a glomerulus *‘i’* and neuron *‘j’* is given by a Poisson distribution with mean=average number of synapses. The average number of synapses is given by the formula: Average number of synapses = number of synapses/ (number of glomeruli * number of neurons).

Here, the number of synapses is the product of the density of synapses in layer 1a and the width of layer1a (Figure 1). Detailed studies on synaptic densities have shown that synaptic densities across various species and brain regions are remarkably similar averaging about a billion synapses per mm^3^ (DeFelipe et al. 2002; Schüz & Palm 1989). By this measure, assuming that layer 1a occupies half the width of layer 1, the number of synapses under 1 mm^2^ is 85 million (since width of layer 1a in the APC is .17 mm, Figure 1). Given 2000 (or 3700) glomeruli and 41,000 neurons under 1 mm^2^ of cortical surface, the mean number of synapses received by a neuron from any glomerulus is around 1 (or .5), as we had previously reported (Srinivasan & Stevens 2013; Srinivasan & Stevens 2012). The second component of connectivity is the synaptic strength for each synapse, which, by estimates in other regions of the brain follows a Gamma distribution (Murthy et al. 1997).

Thus, a model of the feedforward circuit from the bulb to the piriform cortex requires 4 quantities: the number of glomeruli in the bulb, the number of neurons in the piriform cortex, the mean number of synapses between a glomerulus and a neuron, and the strength of each of these synapses. Our report now makes it possible to build such a model by providing the number of piriform neurons, and a way to estimate the synaptic quantities based on our volumetric estimates of Layer 1a and studies of synaptic densities. Further studies to estimate synaptic densities specifically for the piriform cortex would be helpful in developing the accuracy of models even more.

Finally, principal neurons in the piriform cortex receive excitatory and inhibitory inputs from glomeruli (Suzuki & Bekkers 2010; Stokes & Isaacson 2010). The excitatory input takes the form of direct feedforward excitation from M/T cells or glomeruli. The inhibitory input takes the form of indirect inhibition, wherein M/T cells or glomeruli excite inhibitory cells in Layer 1 that then inhibit principal cells. As Stettler and Axel state in their model (Stettler & Axel 2009), net piriform neuron activity must take account of both these inputs and is given by *N* = *Conn_Excite_*. *G* – *Conn_Inhibit_*. *G* (*Conn_Excite_* and *Conn_Inhibit_* are connection matrices between the glomeruli in the bulb and principal cells in the piriform; *Conn_Excite_* and *Conn_Inhibit_* describe the excitatory and inhibitory connections, respectively). The parameters for the inhibitory connections can be estimated by combining our report with a previous study that provided a detailed quantitative description of the inhibitory neurons in the piriform cortex (Suzuki & Bekkers 2010). Notably, this simple model is but a first step in building a model of the piriform cortex that can explain piriform activity (Haberly 2001).

## Methods

The volumetric and neuronal estimates were gathered from 3 C57 adult male mice of age > 8 weeks. Here, we outline our methods for volumetric and neuronal estimates, and start with the gathering of volumetric estimates. The contours of the 3 layers were outlined with Neurolucida (version 10.01; MBF Bioscience) at low magnification (2X and 4X objectives). For each mouse, olfactory regions were outlined based on standard atlases and primary literature (Paxinos & Franklin 2004; Neville & Haberly 2004).

### Alignment

Once the layers were outlined, individual sections were aligned using the following procedure. Starting with the second section, each section was aligned to the previous one. Areas and volumes for the anterior, posterior, and entire piriform cortex were then obtained from 3 dimensional reconstructions with Neurolucida explorer. The surface area of the piriform cortex was estimated with the contours of the outer layer of layer 1. Examples of the 3D reconstructions are show in Suppl. Fig. 1.

### Neuronal density

10- or 20- μm-wide columns, perpendicular to the pial surface, extending down to the boundary of layer 3, were delineated with the Neurolucida contours option. The sections, and the columns within them were chosen so that we had equal coverage along the rostral-caudal and dorsal-ventral extents, and equal amount of coverage of the APC and PPC. For obtaining surface area density counts, three measures were used: the width of the column, the thickness of the section, and the number of neurons in this column. Columns were randomly chosen while ensuring that they were close to perpendicular to our coronal cut and the surface of the brain. Neurons were counted with standard unbiased stereology techniques in Nissl-stained sections at 100X oil magnification (Figure 3). The counting column functioned as a typical dissector whereby neurons and glia were marked as they come into focus given that they also fell within the acceptance lines of the dissector.

**Figure 3:**
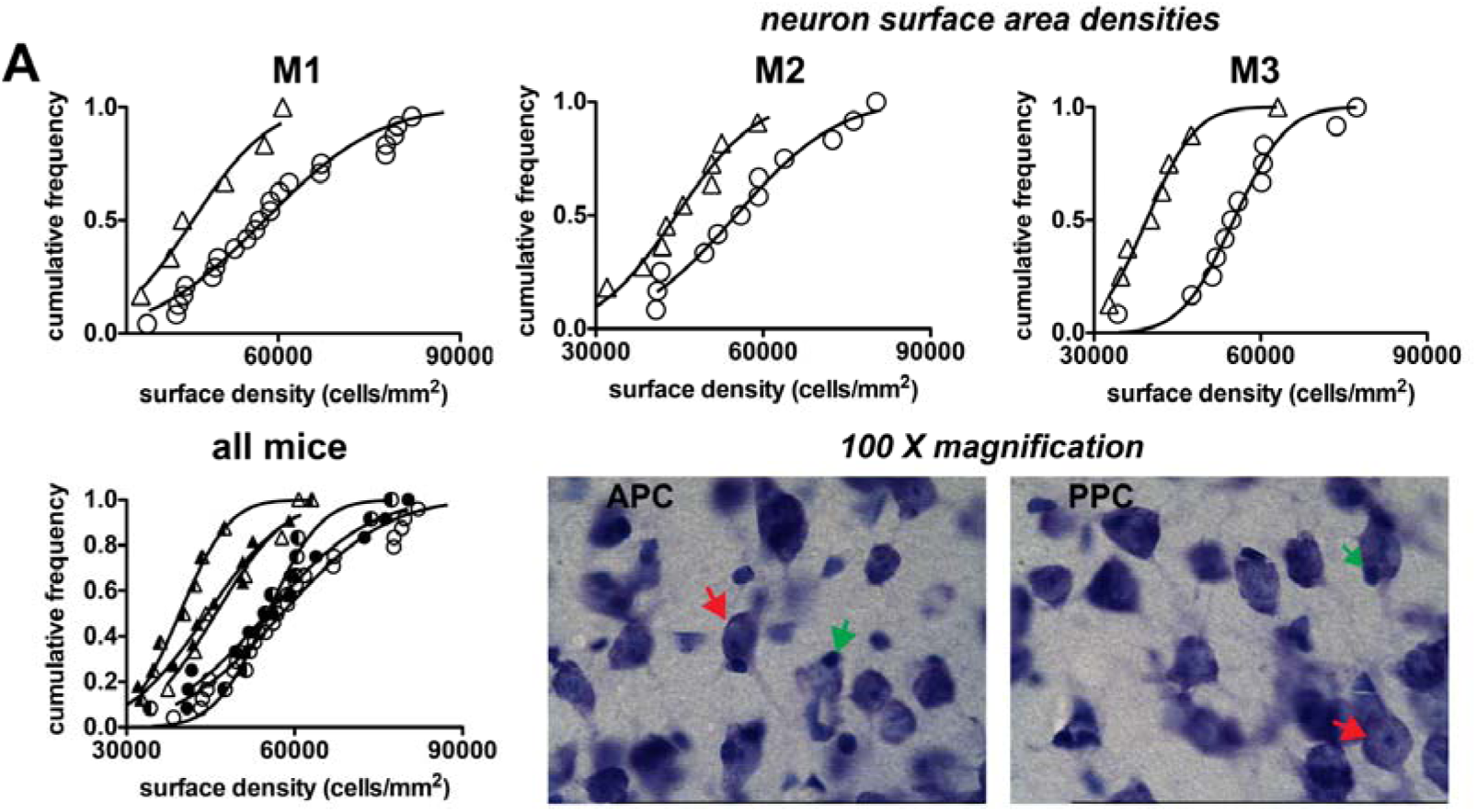
Validation of neuronal surface density counts. (A) Cumulative frequency histograms of density counts in each individual animal for the anterior (circles) and posterior piriform cortex (triangles). Smooth lines are cumulative Gaussians fits with the same mean and standard deviation (R^2^ > .96 for all plots). (B) Samples of APC and PPC sections at 100X magnification, with representative neurons and glia marked by Red and Green arrows.Scale bar:100 μm.

Neurons were differentiated from glia on the basis of size (bigger) and morphology (distinctive shape and processes extending out) of the cell, and by the presence of a nucleolus: glia were also more punctate. A few cells (less than 10 %) were hard to distinguish and were labelled as unknown and not included in the counts. Approximately 50 to 100 objects of interest were counted in each 3D column. The total number of neurons counted ranged from 800 to 1200 neurons per specimen for each region. We counted all the neurons in our column from the pial surface to the white matter without the use of guard zones. A detailed discussion of guard zone use as it pertains to stereological methods and data collection for frozen sections was reported by Carlo and Stevens.

As we used frozen sections, we did not use guard zones (Carlo & Stevens 2011). This brings up an issue of over-counting. We were careful to address this potential issue by marking nucleoli – which typically have a radius of 1.5-2 μm – in 40 μm sections, reducing the margin of error to less than 5 % (Konigsmark 1970; Gramsbergen et al. 1987). We also took care to mark a single nucleolus in those few neurons that had multiple nucleoli (Gramsbergen et al. 1987).

The Neurolucida files for volumetric and neuronal surface area densities are provided as part of the supplement for those interested in a closer analysis.

## Histological Procedures

Brain tissues from the mice were analyzed. Animal care protocols were approved by the Salk Institute Animal and Use Committee and conform to US Department of Agriculture regulations and National Institutes of Health guidelines for humane care and use of laboratory animals. Each specimen was perfused with aldehyde fixative agents and stored long-term in 10% formalin. In preparation for cutting, all brains were submerged in solutions of 10% (wt/vol) glycerol and 10% formalin until the brain sank, and then moved into 20% glycerol and 10% formalin until the brain sank again; the average time in each solution was 3 to 10 d. These cryoprotected brains were then cut coronally on a freezing microtome at a thickness of 40 or 50 μ m. Every 6th section was stained with thionin for visualization of Nissl bodies.

## Thionin Staining

The tissue was defatted with 100% Ethanol: Chloroform (1:1) overnight, rehydrated in a decreasing alcohol (with DI H 2 O) series (100 %, 95, 70, 50), then treated with DI H 2 O, Thionin stain, followed by DI H 2 O, an increasing alcohol series (50 %, 70, 95, 95, 100, 100), Xylenes I, Xylenes II, and then cover-slipped. The tissue was dipped 4-5 times in each of the solutions for 1 minute except for the thionin stain (1-2 minutes), and Xylenes II (1 hour). The Thionin stain was prepared by first gently heating 1428 ml DI H_2_O, 54 ml of 1M NaOH, 18 ml of glacial acetic acid until the solution was steaming. Then, 3.75 gms of Thionin was added and the solution was boiled gently for 45 min, cooled, filtered and used for staining.

## Data Records

The supplement contains 9 Neurolucida files. The first 3 files contain measurements for calculating neuronal densities. Neurons were marked with a + symbol. The next 6 files contain measurements from aligned files for estimating surface areas and volumes using Neurolucida explorer. The suffix msX indicates the mouse number (X = 1,2, or 3).

Counts of neurons for individual columns can be obtained by opening the measurement files in Neurolucida. Volumetric estimates can be obtained by opening the aligned files in Neurolucida explorer and using the statistical descriptor tool to obtain estimates.

## Technical Validation

The neuronal density counts were validated by cumulative frequency Gaussians shown in Figure 3. The spatial distribution of neurons within a region can take two forms. It can have constant density or be distributed non-homogeneously. In the case of constant density, because biology is variable, neurons will have a uniform density across the region but will be distributed according to some random process at smaller scales. In most cases this distribution is Poisson, i.e. the probability of encountering a neuron as one traverses a straight line is given by a Poisson. In the limit, the Poisson distribution can be approximated by a Gaussian or Normal distribution.

What this means for our study is that the neuronal densities for all columns within the same mouse should also follow a Gaussian, and provide a way to test the validity of the counts. This is exactly what we observed, when we plotted the cumulative frequency histogram and compared it with a cumulative Gaussian (Figure 3).

## Acknowledgements

We would like to thank the Kavli Institute at UCSD and the National Science Foundation (NSF) directorate of Mathematical and Physical Sciences for their support through the 2014 NSF Early-Concept Grants for Exploratory Research (NSF PHY-1444273) for their support, and Chris Kintner and Jeffry Isaacson for comments on the project.

## Author contributions

SS performed the experiments, SS and CFS conceived and developed the project, SS and CFS wrote the report.

